# Arabidopsis CPR5 regulates ethylene signaling via interacting with ETR1 N-terminus and controlling mRNAs nucleocytoplasmic transport

**DOI:** 10.1101/862862

**Authors:** Jiacai Chen, Yanchong Yu, Xinying Sui, Longfei Qiao, Chun-Hai Dong

**Affiliations:** College of Life Sciences, Qingdao Agricultural University, Qingdao 266109, China

**Keywords:** CPR5, ethylene, nucleoporin, nucleocytoplasmic transport, Arabidopsis

## Abstract

ETR1 is the major ethylene receptor in *Arabidopsis thaliana*. Previous studies showed that RTE1 and CPR5 can bind to ETR1 and play regulatory roles in ethylene signaling. RTE1 has been suggested to promote ETR1 signal transduction by influencing the conformation of ETR1, but little is known about the mechanism of CPR5 on the regulation of ETR1 signaling. In this study, we showed that both CPR5 and RTE1 could interact with the N-terminal transmembrane domains of ETR1, and CPR5 needs at least three transmembrane domains of ETR1 while RTE1 needs only two for the binding. As CPR5 has also been shown to be localized in the nuclear membrane and might act as a nucleoporin, we analyzed the effects of CPR5 on the nucleocytoplasmic transport of ethylene-related mRNAs using poly(A)-mRNA *in situ* hybridization and real-time quantitative PCR (qPCR), and the results indicated that CPR5 could selectively regulate the nucleocytoplasmic transport of mRNAs in ethylene signaling pathway. In contrast, the nucleoporin mutants (*nup160*, *nup96-1* and *nup96-2*) dramatically accumulated all the examined mRNAs in the nucleus. In conclusion, the present study provides evidence demonstrating that CPR5 regulates ethylene signaling through interacting with the ETR1 receptor and controlling the mRNAs nucleocytoplasmic transport in ethylene signaling pathway.

**Key message:** This study reveals that CPR5 is involved in the regulation of ethylene signaling via two different ways: interacting with the N-terminal domains of ERT1 and controlling the nucleocytoplasmic transport of mRNAs in ethylene signaling pathway.

## Introduction

Ethylene is the simplest gaseous plant hormone and widely distributed in plant tissues and cells. Previous studies have shown that ethylene, as an important plant hormone, plays important roles in regulating plant growth and development, such as seed germination, fruit maturity, flower development, sex determination (Kamachi *et al.*, 1997; Yamasaki *et al.*, 2000), and leaf development, senescence and abscission (Bieleski and Reid, 1992; Kieber and Ecker, 1993; Guerrero *et al.*, 1998). Ethylene is also involved in plant responses to biotic and abiotic stresses, including resistance to hypoxia (Fukao and Bailey-Serres, 2008; Guillaume and Margret, 2008; Magneschi and Perata, 2009; Justin and Armstrong, 2010), drought and salt stresses, inhibition of hypocotyl elongation in seedlings under dark conditions, promotion of hypocotyl growth under light conditions, and relief of photooxidation stress (Roman and Ecker, 1995; Bleecker and Kende, 2000). When ethylene or ethylene precursor ACC was added to the medium, the dark-grown Arabidopsis seedlings exhibited typical characteristics, termed the ethylene “triple response” with exaggerated apical hook, inhibited root and hypocotyl elongation, and swelled hypocotyl (Bleecker *et al.*, 1988; Guzmán and Ecker, 1990). According to the ethylene “triple response”, a large number of Arabidopsis mutants with altered ethylene sensitivity were isolated. Based on the genetic study of the ethylene responsive mutants, a linear ethylene signal transduction pathway emerged (Bleecker *et al.*, 1988; Guzmán and Ecker, 1990; Kieber *et al.*, 1993; Roman and Ecker, 1995; Guo and Ecker, 2004).

Ethylene signaling pathway starts from the binding of ethylene with receptors, and then through the transmission of downstream factors, and finally reaches the nucleus and activates the expressions of the ethylene responsive factors, thereby resulting in various ethylene responses. So far, many components and regulatory factors in the pathway have been isolated, such as ethylene receptor family (ETR1, ETR2, ERS1, ERS2, EIN4) (Chang *et al.*, 1993; Hua *et al.*, 1998; Hua and Meyerowitz, 1998; Sakai *et al.*, 1998), an MAPKKK protein CTR1 (Kieber *et al.*, 1993), the endoplasmic reticulum (ER) membrane-localized protein EIN2 (Alonso *et al.*, 1999), the transcription factor EIN3 and EIN3-LIKE1 (EIL1) (Chao *et al.*, 1997; Solano *et al.*, 1998), and the downstream ethylene-responsive factors (ERFs) (Fujimoto *et al.*,2000).

Ethylene receptors mainly exist in ER and Golgi in the form of dimer, and its N-terminal hydrophobic region needs copper ion as cofactor when binding to ethylene (Chen *et al.*, 2002; Schaller and Bleecker, 1995; Rodriguez *et al.*, 1999; Binder *et al.*, 2010). Among the five ethylene receptors in Arabidopsis, the ETR1 was believed to play a major role in ethylene signaling. To explore the regulatory mechanism of the ETR1 receptor, we and colleagues recently reported the isolation of the ETR1 receptor-associated protein RTE1 and CPR5 based on their regulatory functions in the ETR1 receptor signaling (Resnick *et al.*, 2006; Zhou *et al.*, 2007; Dong *et al.*, 2008; Dong *et al.*, 2010; Wang *et al.*, 2017).

Arabidopsis *RTE1* gene encodes a protein containing 250 amino acids which belongs to a membrane protein. By co-localization with ETR1, it is mainly localized in ER and Golgi (Zhou *et al.*, 2007; Dong *et al.*, 2008). There are three homologues of *RTE1* in tomato, one of which has been proved to be involved in the regulation of ethylene response in crops (Barry and Giovannoni, 2006; Harry, 2006; Ma *et al.*, 2012). The rice *RTE1* homologues *OsRTH1* and *RTE*-like genes (*DeRTE1* and *DeRTH1*) in carnation have also been shown to be involved in the regulation of ethylene effect on seedling growth and flower senescence (Yu *et al.*, 2011; Zhang *et al.*, 2012).

RTE1 is a positive regulator of ethylene receptor ETR1 and can interact directly with ETR1 (Resnick *et al.*, 2006; Dong *et al.*, 2010). Genetic analysis showed that RTE1 is essential for ETR1 to function in Arabidopsis, but not for other ethylene receptors (Resnick *et al.*, 2006; Resnick *et al.*, 2008). Studies have shown that RTE1 might function via influencing the conformation of ETR1 receptor. The target site of RTE1 on ETR1 might be the ethylene binding domain on ETR1, thus it was suggested that RTE1 promoted ETR1 signal transduction by influencing the conformation of the ethylene binding domain of ETR1 (Resnick *et al.*, 2008).

CPR5 is initially isolated from the research on plant systemic acquired resistance (Bowling *et al.*, 1997; Boch *et al.*, 1998). Studies revealed that CPR5 participates in the regulation of different physiological and pathological processes in plants, such as K^+^ dynamic balance, ABA signal transduction, redox balance, programmed cell death, and ROS status and signal transduction (Kirik *et al.*, 2001; Jing *et al.*, 2005; Jing and Dijkwel, 2008; Jing H *et al.*, 2010). CPR5 is also involved in the regulation of gene replication, cell division, cell proliferation and spontaneous cell death (Brininstool *et al.*, 2008; Perazza *et al.*, 2011). It was suggested that CPR5 is one of the cell cycle regulators and plays a key role in this process (Bao and Hua, 2014). Recent studies have shown that CPR5, as a nucleoporin, plays a role in controlling of triggering immunity and programmed cell death (Gu *et al.*, 2016). The regulatory function of CPR5 in ethylene signaling was also reported (Aki *et al.*, 2007; Jing *et al.*, 2007; Wang *et al.*, 2017). The seedlings of Arabidopsis *cpr5*/*hys1*/*old1* mutants exhibited enhanced ethylene sensitive phenotype with the addition of ethylene precursor ACC. Intrigually, it was found that either overexpressed or knockout of *CPR5* could lead to increase of the ethylene sensitivity (Wang *et al.*, 2017), suggesting that CPR5 might act in a different manner from RTE1 in regulating the ETR1 receptor signaling.

Subcellular localization analysis indicated that CPR5 is localized in the endometrial system (ER and Golgi) and nuclear membrane (Gu *et al.*, 2016; Wang *et al.*, 2017). CPR5 was shown to be able to directly interact with the ETR1 receptor (Wang *et al.*, 2017), providing a clue that it might compete with RTE1 to bind to the ETR1 receptor and affecting ethylene signaling. Meanwhile, CPR5 might aslo act as a nucleoporin in controlling the nucleocytoplasmic transport of mRNAs in ethylene signaling pathway.

In this study, we used different approaches to investigate the regulatory mechanism of CPR5 in ethylene signaling pathway. Genetic analysis provided evidence supporting that the mode of CPR5 regulating the ETR1 receptor signaling could be different from that of RTE1. By yeast split-ubiquitin and bimolecular fluorescence complementation (BiFC) assay, we showed that both CPR5 and RTE1 could interact with the N-terminal transmembrane domains of ETR1, and it is likely that there might exist a competitive binding between CPR5 and RTE1 when they interact with ETR1. Using poly(A)-mRNA *in situ* hybridization and real-time quantitative PCR, we analyzed the effects of CPR5 on the nucleocytoplasmic transport of ethylene signaling related mRNAs, and the results indicated that CPR5 could selectively regulate the nucleocytoplasmic transport of mRNAs in ethylene signaling pathway. As controls, the nucleocytoplasmic transport of all the examined mRNAs in the nucleoporin mutant *nup160, nup96-1*, and *nup96-2* were affected. These observations significantly advanced our understanding of the regulatory mechanism of CPR5 in ethylene signaling pathway.

## Materials and Methods

### Plant materials and and ethylene response assays

Seeds of the wild type (WT, Col-0) or mutants were surface sterilized and then sowed on 1/2 MS (Murashige and Skoog) medium or soil in a controlled environment growth chamber set at 21°C under 16 h light/8 h dark. The Arabidopsis ethylene response mutants were ether obtained from previous studies, or generated by genetic crossing (Dong *et al.*, 2008; Wang *et al.*, 2017; Zheng *et al.*, 2017). The F2 progeny from the crosses were screened by specific PCR markers, as previously described (Resnick *et al.*, 2006; Wang *et al.*, 2017). The Arabidopsis nucleoporin mutant *nup160* was as previously described (Dong *et al.*, 2006). The mutants *nup96-1* and *nup96-2* were obtained from Dr. Xin Li (Zhang and Li, 2005).

The ethylene response of Arabidopsis seedlings was examined in the presence of ACC at different concentrations (0, 0.5, 5, 10, 20, or 100 μM) as previously described (Wang *et al.*, 2017). After sterilization, seeds were sowed on 1/2 MS medium containing ACC, and treated at 4°C for 3 d. Thereafter, the plates were moved to a growth chamber for 8 h under white light, and then, the plates were wrapped with aluminum foil and placed in a growth chamber for the indicated periods. Measurement of hypocotyl length and statistical data were evaluated either by Student’s *t*-test or Fisscher’s test for a multiple comparison.

### Yeast split-ubiquitin assay

For yeast split-ubiquitin assay, the portions of Arabidopsis ETR1 (1-368 aa; 369-596 aa; 597-739 aa; 1-78 aa; 79-139 aa and 140-368 aa) were each PCR-amplified from an existing template, cloned into PMD18-T vector and then transferred into a bait vector pPR3-N or a prey vector pBT3-STE through the restriction sites shown in supplementary table (Table S1). The final constructs were verified by sequencing of the inserts. The bait or prey vectors harboring a full-length of *ETR1*, *CPR5* or *RTE1* were obtained from a previous study (Wang *et al.*, 2017).

### Bimolecular fluorescence complementation (BiFC) assay

To generate constructs harboring the fusions of *cYFP-ETR1* (1-368 aa) and *cYFP-ETR1* (1-78 aa), the coding sequences of *cYFP* (466-717 aa), *ETR1* (1-368 aa) and *ETR1* (1-78 aa) were each PCR-amplified from existing templates as previously described (Wang *et al.*, 2016), and cloned into the pMD18-T vector. After sequencing verification of the inserts, the fragments were cloned into a binary vector pCambia1300-3HA through restriction sites (Xba I and Kpn I for *cYFP*; Kpn I and BamH I for *ETR1* (1-368 aa) and *ETR1* (1-78 aa). The contructs containing the fusions of *nYFP-CPR5* and *nYFP-RTE1* were obtained from a previous study (Wang *et al.*, 2017). Transformation mediated by Agrobacterium (GV3101) using onion epidermal cells was accoding to a previous study (Xu *et al.*, 2014). Fluorescent signal from the transfected cells was examined under a fluorescence microscope (Nikon ECLIPSE Ti-S, Japan) or a laser scanning confocal microscope (Leica TCS SP5, Germany). The primers used for the constructs of BiFC assay are listed in supplementary table (Table S2).

### RNA extraction and real-time quantitative PCR analyses

Total RNA was extracted from Arabidopsis seedlings according to TRNzol (TIANGEN, China). Half of the same seedlings were used to extract nuclear RNA. 3 g of fresh seedlings were ground into powder in liquid nitrogen, and 6 mL extract A (0.3 M sucrose, 50 mM Tris-HCl, pH 8.5) was added to the ground material. After further grinding, the residue was filtered by miracloth, and the filtrate was collected in the RNase free tube. The residue was added to the extract A to continue grinding and filtering. The two filtrates were mixed and centrifuged at 500 g under 4°C for 5 min. The supernatant was discarded, and then 1 mL TRNzol reagent was added to the sediment, and RNA was extracted as instructed.

cDNA was synthesized with oligo(dT) primers using PrimeScript™ 1st strand cDNA synthesis kit (Takara, Japan). Quantitative RT-PCR analysis was performed on an Agilent Real-Time qPCR apparatus (Mx3000P system) using SYBR Premix ExTaq TM II (Takara, Japan). Biological replicates for each set of experiments were carried out three times, and the mean value of three replicates was normalized using *Tubulin8* as an internal control. Quantitative PCR was conducted at 95°C for 10 min, followed by 40 cycles at 95°C for 30 s, 55°C for 30 s, and 72°C for 20 s. The primers used for quantitative RT-PCR are listed in supplementary table (Table S3).

### Poly(A)-mRNA in situ hybidization

Examination of the Poly(A)-mRNA *in situ* localization was performed as previously described (Gong *et al.*, 2005). In brief, samples were taken from equivalent portions of young leaves of Arabidopsis plants grown for 3 weeks. After fixation, dehydration and prehybridization, the samples were hybridized with the oligo-dT labeled with 6-FAM fluorescein at N-terminus (synthesized by Sangon Biotech, China) at 50°C in darkness for more than 8 h. After washing, the samples were observed immediately using a laser scanning confocal microscopy (Leica TCS SP5, Germany) with a 488-nm excitation laser. Each experiment was repeated at least three times, and similar results were obtained.

## Results

### CPR5 could bind to the N-terminal domains of ETR1 receptor

As both CPR5 and RTE1 could directly interact with ETR1 and act as the regulators of the ETR1 signaling, we are interested to know whether the CPR5 and RTE1 could bind to the same region on the ETR1 receptor. To give insight into this question, we first investigated the binding sites of CPR5 and RTE1 to the ETR1 receptor, respectively. The ETR1 was divided into three segments with different domains: ETR1 (1-368 aa) including three transmembrane domains and the GAF domain, ETR1 (308-596 aa) containing the histidine kinase domain, and ETR1 (604-739 aa) containing the response regulatory domain (Fig. 1A). Yeast split-ubiquitin assays showed that both CPR5 and RTE1 could only interact with the ETR1 (1-368 aa), but not the other two regions (Fig. 1B, C).

**Fig1.**
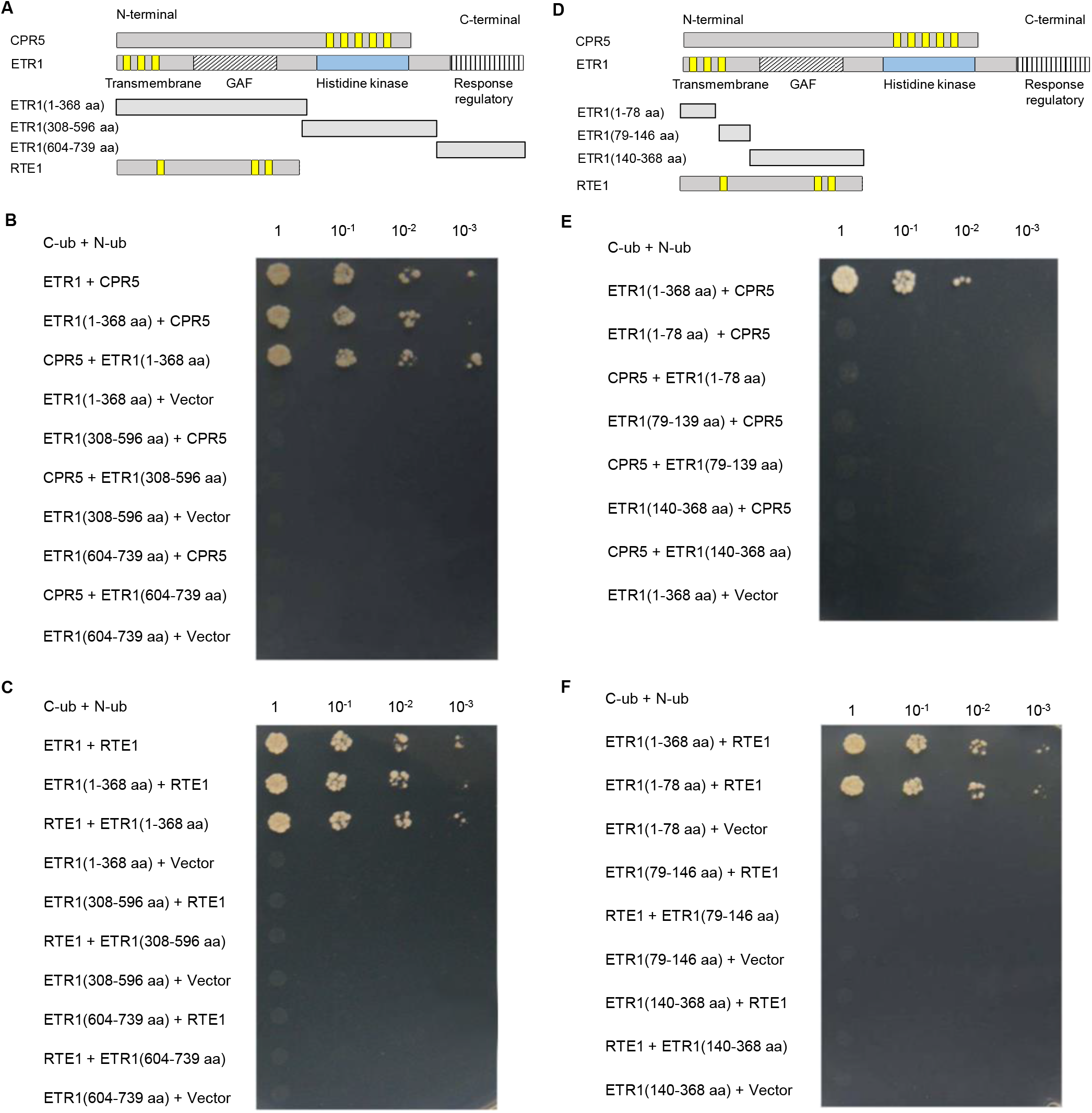
Analysis of the binding sites of CPR5 and RTE1 to ETR1 by yeast split-ubiquitin assay. (A & D) Diagram for the full length of CPR5, ETR1 and RTE1, and the truncated portions of ETR1. (B, C, E & F) Molecular interaction of CPR5 (B & E) and RTE1 (C & F) with the portions of ETR1 in the yeast split-ubiquitin assay. The bait protein CPR5 (B & E) and RTE1 (C & F) was paired with the full length or portions of ETR1. Positive interaction is indicated by growth on medium lacking leucine, tryptophan, histidine and alanine (-LTHA). Undiluted and diluted liquid cultures were spotted on the plates and incubated for 5 d at 30°C.

In order to know the precise domain in which CPR5 and RTE1 could directly interact with ETR1, we further divided the ETR1 (1-368 aa) into three segments: ETR1 (1-78 aa) containing two transmembrane domains, ETR1 (79-146 aa) containing one transmembrane domain, and ETR1 (140-368 aa) containing the GAF domain only (Fig. 1D). The yeast split-ubiquitin assays showed that CPR5 did not interact with any of the three segments, while RTE1 could interact with the ETR1 (1-78 aa) segment, but not the other two segments (Fig. 1E, F).

Further evidances were given by the bimolecular fluorescence complementation (BiFC) assay. The BiFC assay results showed that CPR5 could interact with the ETR1 (1-368 aa), but not the ETR1 (1-78aa), whereas RTE1 was able to interact with the ETR1 (1-78aa) (Fig. 2), being consistent with the above results.

**Fig. 2.**
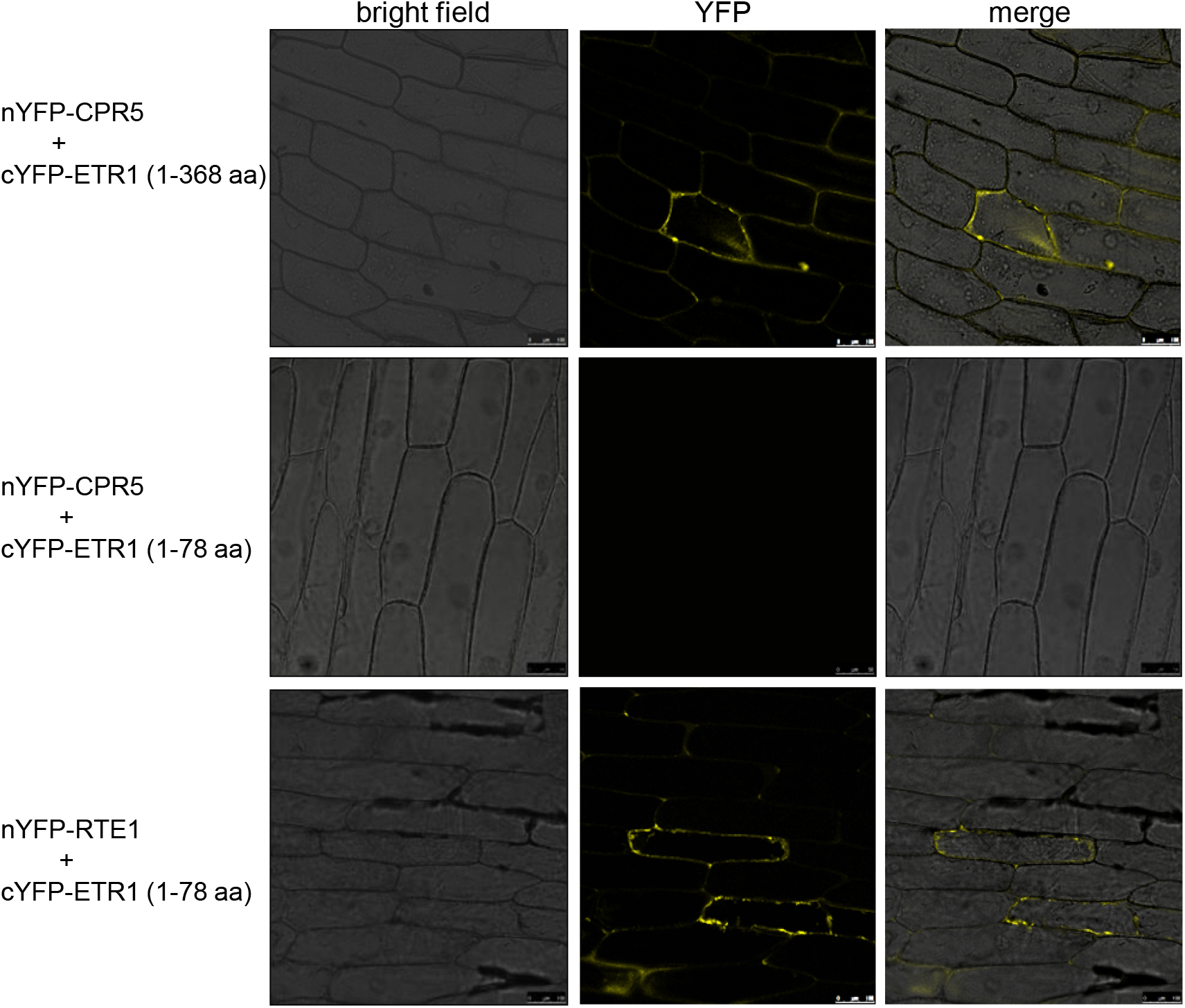
BiFC assay for the molecular association of CPR5, RTE1 with the ETR1 portions in onion epidermal cells. The N-terminal half of YFP (nYFP) was fused to CPR5 or RTE1, and the C-terminal half of YFP (cYFP) was fused to ETR1 (1-368 aa) or ETR1 (1-78 aa), respectively, and the constructs were co-infiltrated into onion epidermal cells. Fluorescent signal of YFP was detected by laser scanning confocal microscopy at 505-530 nm. Bars = 50 μm.

In conclusion, both CPR5 and RTE1 could bind to the N-terminus of ETR1, the RTE1 could bind to a smaller region comtaining two transmembrane domains, while the CPR5 binding required at least three transmembrane domains and the GAF domain of ETR1. Although the region sizes of ETR1 required for the interactions with CPR5 and RTE1 was different, the binding sites were generally the same, it was speculated that there might exist a competitive binding between CPR5 and RTE1 when they interacted with the ETR1 receptor.

### CPR5 might act downstream of RTE1 and play a more basic regulatory function in ethylene signaling

As knockout of *CPR5* or *RTE1* could restore the ethylene sensitivity of *etr1-2* (Resnick *et al.*, 2006; Wang *et al.*, 2017), it is interesting to see if additative effect exists on the ethylene sensitivity of *etr1-2* when both CPR5 and RTE1 were knocked out from the mutant. The double mutant of *cpr5-T3 rte1-3* was generated by a genetic cross between *cpr5-T3* and *rte1-3*, and the ethylene “triple response” analysis was carried out. It was shown that the hypocotyl lengths were of the most significant difference between the WT and mutants under low concentrations of ACC (0.5, 5 μM) (Fig. 3A, B). With the increase of ACC concentration, the seedling growth of *cpr5-T3* and *cpr5-T3 rte1-3* decreased most, and the hypocotyl lengths of the *cpr5-T3* and *cpr5-T3 rte1-3* were shorter than that of *rte1-3*, indicating that knockout of *CPR5* had stronger effect than that of *RTE1* on the ethylene sensitivity. In addition, the hypocotyl length of *cpr5-T3 rte1-3* was closer to that of *cpr5-T3* when treated with different concentrations of ACC, suggesting that CPR5 might act downstream of RTE1 and play a more general regulatory function in ethylene signaling.

**Fig. 3.**
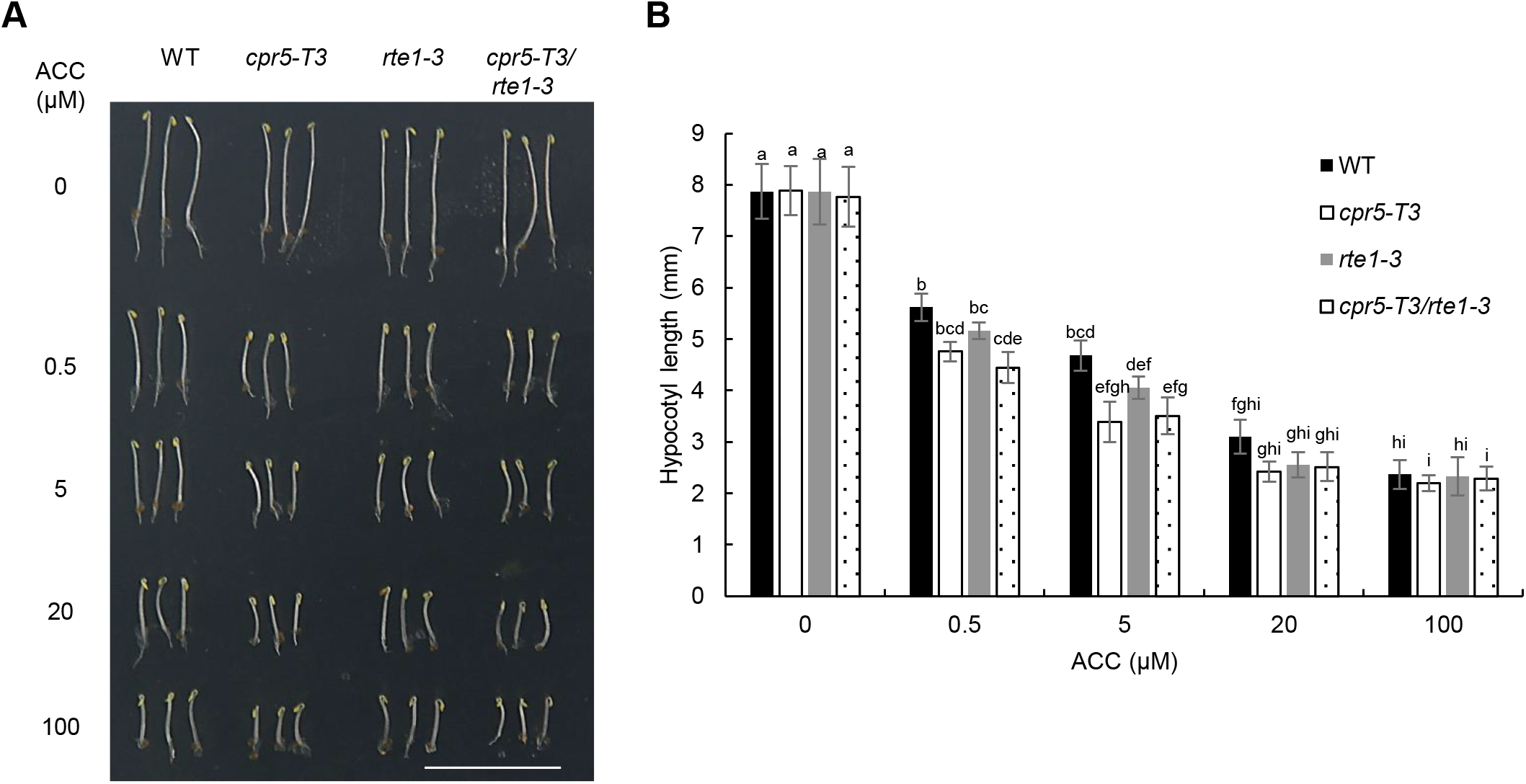
Ethylene “triple response” assays in double mutants. (A) Ethylene sensitivity analysis of the WT (Col-0) and mutants treated with or without ACC for 4 d. Bar= 10 mm. (B) Quantitative analysis of hypocotyl lengths for the wild type (WT) and mutants on 1/2 MS medium with different concentrations of ACC. Significant differences between measurements (P<0.05) are indicated by different letters above the error bars (n=25).

### CPR5 functions in controlling the nucleocytoplasmic transport of ETR1 mRNA

As CPR5 could act as a nucleoporin in controlling of triggering immunity and programmed cell death (Gu *et al.*, 2016), we examined whether CPR5 could function in controlling the nucleocytoplasmic transport of mRNAs in ethylene signaling pathway. Using *in situ* poly(A)-mRNA hybridization, we showed that there was obvious fluorescence aggregation in the nucleus of *cpr5-T3*, while the fluorescence in WT, *rte1-3* and *rth-1* cells was dispersed (Fig. 4). The results indicated that the deficiency of CPR5 could hinder the transfer of mRNAs from nucleus to cytoplasm, thereby resulting in the accumulation of mRNAs in nucleus.

**Fig. 4.**
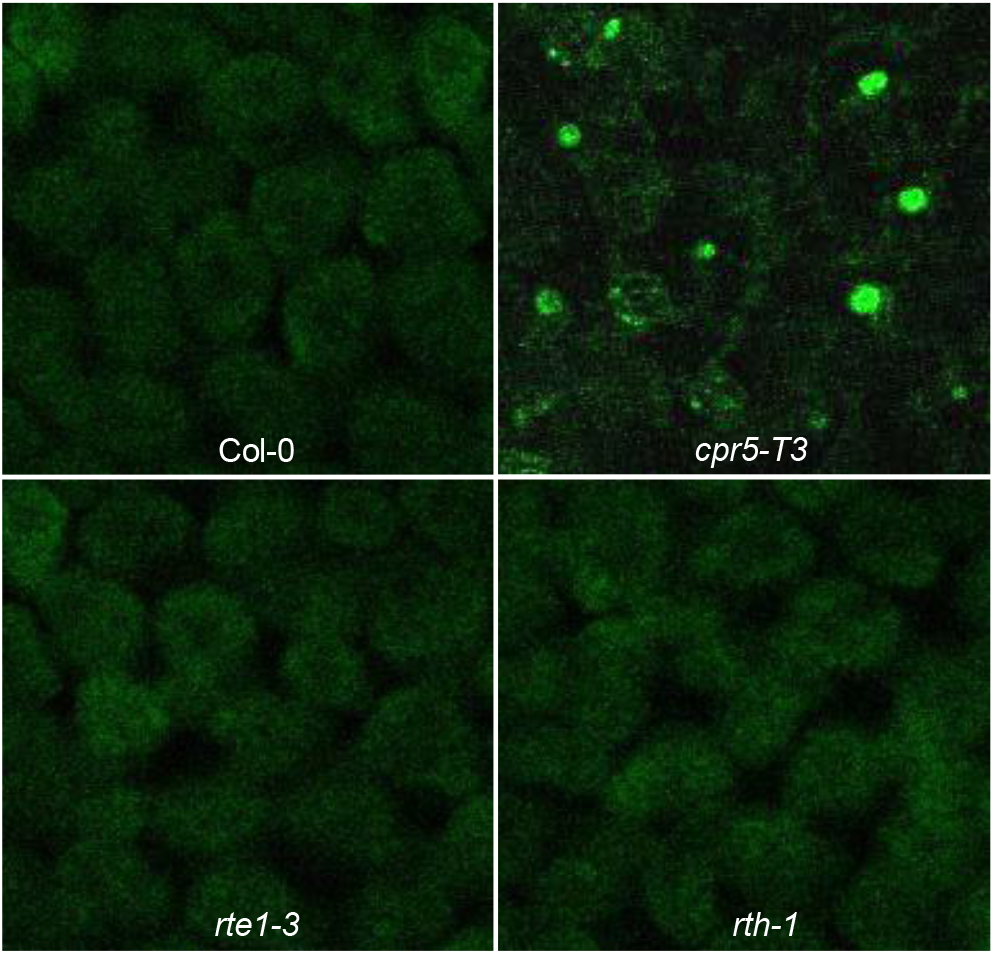
Defect of the cpr5-T3 mutant plants in mRNA export. Poly(A)-mRNA export is blocked in the *cpr5-T3* mutant cells. WT (Col-0), *cpr5-T3, rte1-3* and *rth-1* mutant plants were grown at 22oC for 3 weeks. *In situ* hybridization with fluorescein-labeled oligo (dT) probe was performed using young leaf tissues. Fluorescent signal was examined by a laser scanning confocal microscopy at emission 505-530 nm.

Next, we analyzed the nuclear accumulation of the *ETR1* mRNA. Total RNA and the nucleus RNA were respectively extracted from the 10-day-old seedlings of WT, *cpr5-T3*, *rte1-3* and *rth-1*. qRT-PCR analysis showed that the accumulation of the *ETR1* mRNA in the nucleus of *cpr5-T3* accounted for more than 60% of its total level, while those in the nucleus of WT, *rte1-3* and *rth-1* accounted for about 30%, only half of its accumulation in the nucleus of *cpr5-T3*. As a control, the *Actin1* mRNA accumulation in the nucleus of *cpr5-T3* was almost the same as in WT, *rte1-3* and *rth-1* (Fig. 5). Obviously, these results indicated that CPR5 could seriously affect the nucleocytoplasmic transport of the *ETR1* mRNA.

**Fig. 5.**
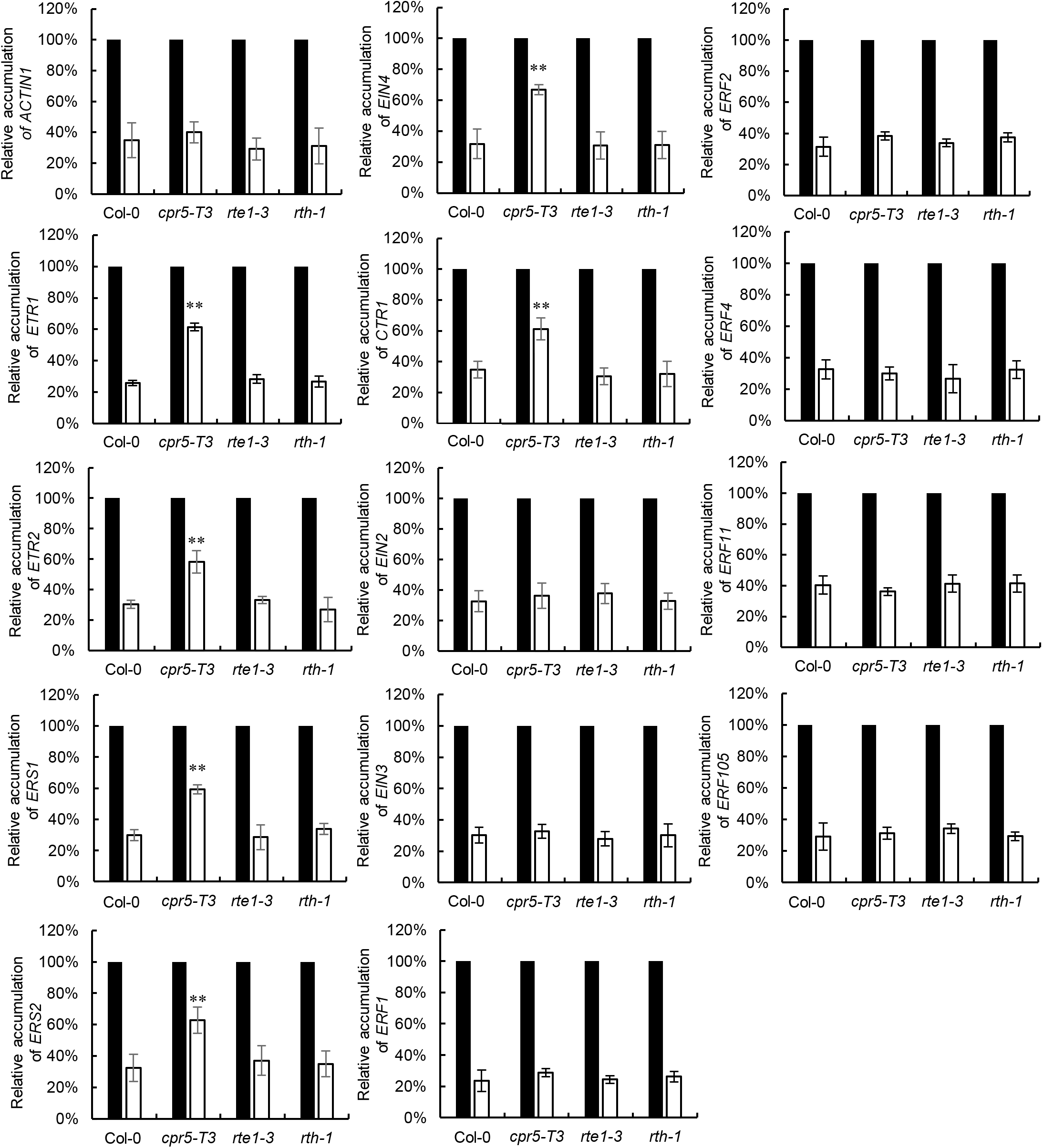
Analysis of the relative accumulation of mRNAs in the nucleus of WT (Col-0), *cpr5-T3, rte1-3* or *rth-1* by qPCR. Plants grown on 1/2 MS medium for 10 d were used for RNA extract. Relative accumulation of nuclear RNA/total RNA was calculated based on qPCR measurements. Black column, total RNA; white column, nucleus RNA. Values are mean ±SD; **p<0.01.

### Nucleocytoplasmic transport of the ethylene signaling related mRNAs was affected in cpr5-T3

We further examined the nucleocytoplasmic transport of the other ethylene signaling related mRNAs in the *cpr5-T3*, including the other ethylene receptors encoding genes (*ETR2*, *ERS1*, *ERS2* and *EIN4*), the downstream key components (*CTR1*, *EIN2*, *EIN3*), and some of the ethylene-induced *ERFs* (Fig. 5). Interestingly, it was observed that the mRNAs of the *ETR2*, *ERS1*, *ERS2*, *EIN4* and *CTR1* were significantly accumulated in the nucleus of *cpr5-T3*, and their accumulation levels were more than 60%, being similar to that of the *ETR1*. However, the mRNAs of *EIN2*, *EIN3*, and the *ERFs* (*ERF1*, ERF2, *ERF4*, *ERF11*, *ERF105*) did not dramatically accumulate in the nucleus of *cpr5-T3* compared to the WT. In contrast, there was no abnormal mRNAs accumulation in the nucleus of *rte1-3* and *rth-1*, indicating that RTE1 and its homolog (RTH) did not act as CPR5 in controlling the nucleocytoplasmic transport of mRNAs.

### Bulk mRNA nucleocytoplasmic export was defected in the nucleoporin mutants (nup160, nup96-1 and nup96-2)

In order to know whether CPR5 could function in controlling the nucleocytoplasmic transport of mRNAs as the other nucleoporins, we analyzed the the nucleocytoplasmic transport of mRNAs in the nucleoporin mutants *nup160*, *nup96-1* and *nup96-2*. *In situ* poly(A)-mRNA hybridization was employed as above. The results revealed that there was obvious fluorescence aggregation in the nucleus of *nup160*, *nup96-1* and *nup96-2*, but no such strong signal in WT (Fig. 6), indicating that a large amount of RNA was accumuled in the nucleus of *nup160*, *nup96-1* and *nup96-2*.

**Fig. 6.**
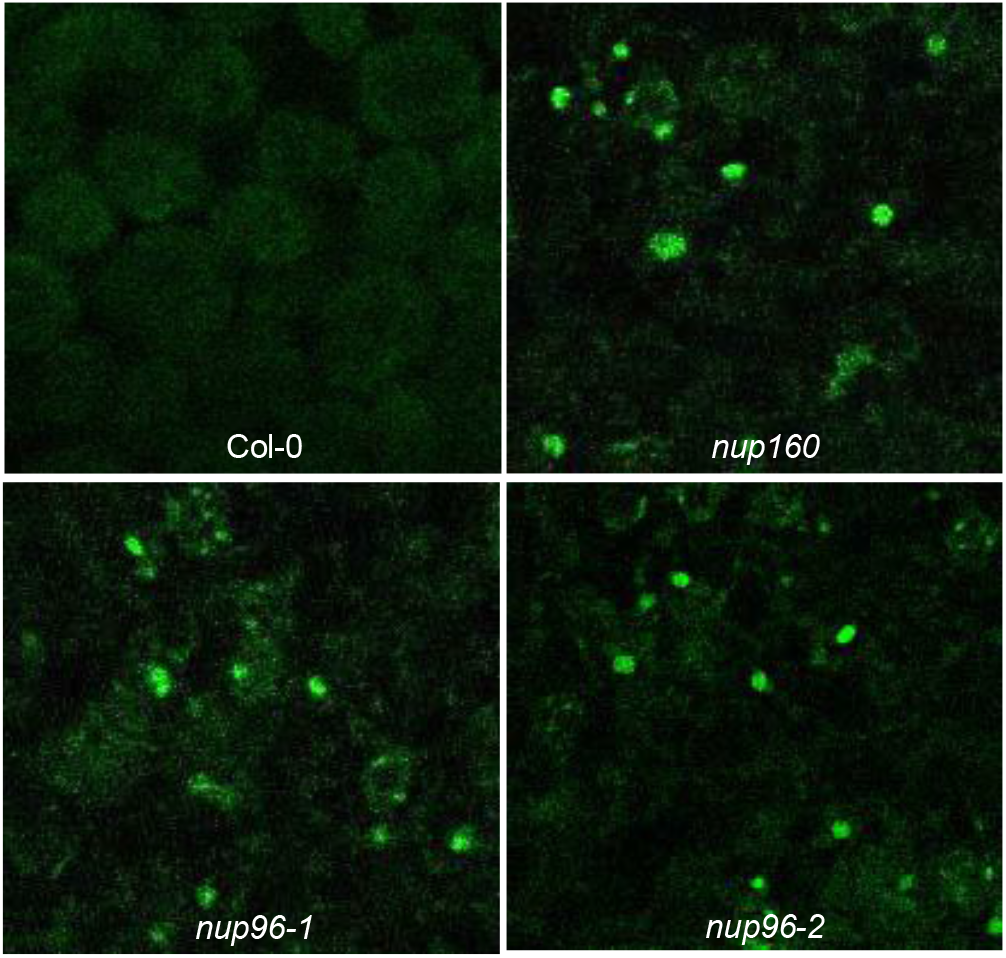
Defect of the *nup160, nup96-1* and *nup96-2* mutant plants in mRNA export. Poly(A)-mRNA export is blocked in the *nup160, nup96-1* and *nup96-2* mutant cells. Col-0 and *nup160, nup96-1* and *nup96-2* plants were grown at 22°C for 3 weeks. *In situ* hybridization with fluorescein-labeled oligo (dT) probe was performed using young leaves. Fluoresecent signal was examined by a laser scanning confocal microscopy (Leica TCS SP5) at emission 505-530 nm.

We next analyzed the relative accumulation of the ethylene signaling related mRNAs in *nup160*, *nup96-1* and *nup96-2*. As shown in fig. 7, the accumulation of all the examined ethylene signaling related mRNAs in the nucleus of *nup160*, *nup96-1* and *nup96-2* reached more than 50% of their total levels, while the accumulation of mRNAs in WT was only about 30%. These results indicated that bulk mRNA cytoplasmic export including all the examined mRNAs was defected in the nucleoporin mutants (*nup160*, *nup96-1* and *nup96-2*).

**Fig. 7.**
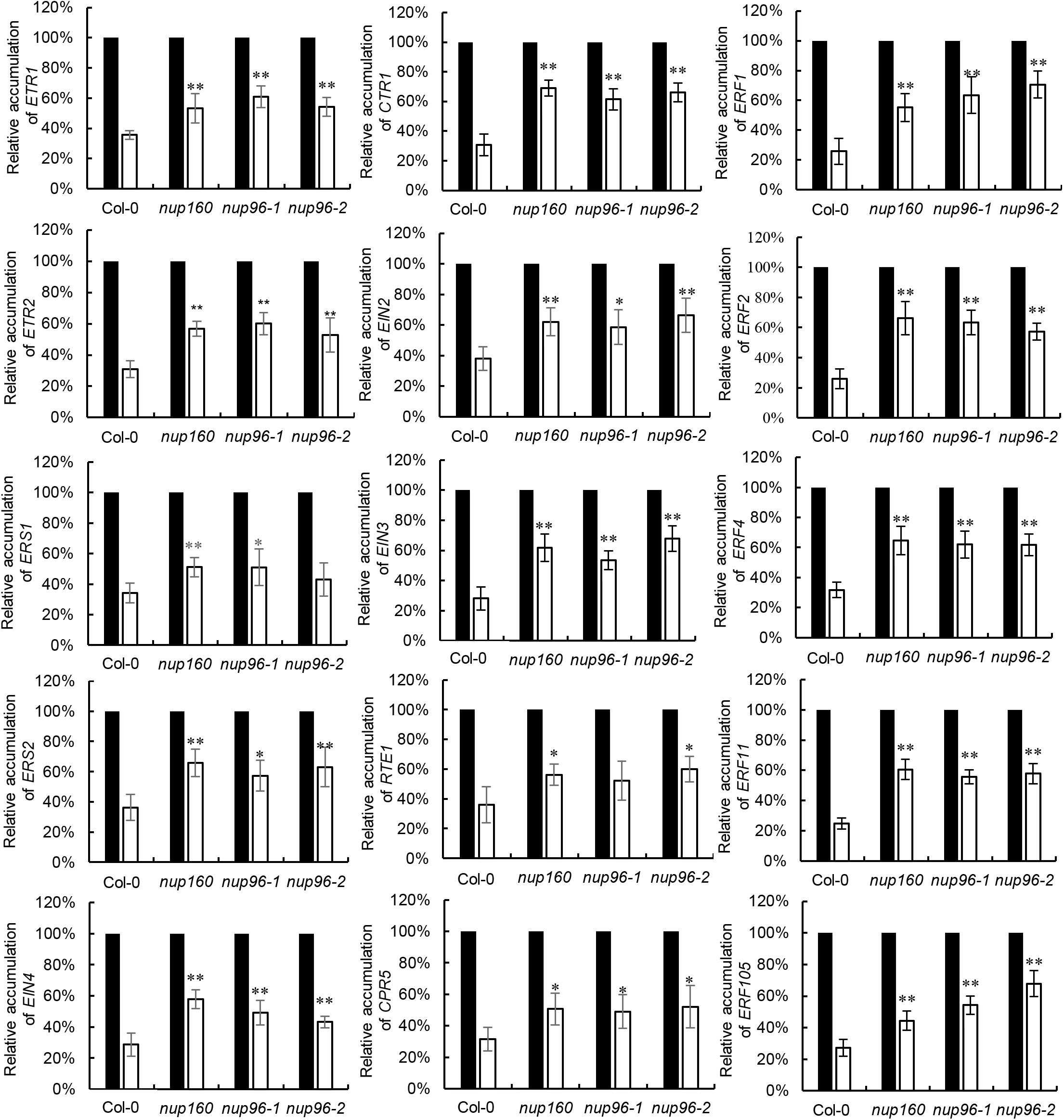
Analysis of the relative accumulation of mRNAs in the nucleus of WT (Col-0), *nup160, nup96-1* or *nup96-2* by qPCR. Plants grown on 1/2MS medium for 10 d were used for RNA extract. Relative accumulation of
nuclear RNA was calculated based on qPCR measurements when total RNA was as one unit. Black column, total RNA; white column, nucleus RNA. Values are mean ± SD; *p<0.05, **p<0.01.

### Ethylene sensitivity was enhanced in nup160, nup96-1 and nup96-2

Defective nucleocytoplasmic transportation of bulk mRNAs in *nup160*, *nup96-1* and *nup96-2* prompted us to investigate the ethylene sensivity of these mutants. When treated with different concentrations of ACC, the hypocotyl lengths of *nup160*, *nup96-1* and *nup96-2* were significantly shorter than that of WT (Fig. 8A, B), and the relative changes in hypocotyl lengths indicated that the nucleoporin mutants *nup160*, *nup96-1* and *nup96-2* were more sensitive to ACC than the WT.

**Fig. 8.**
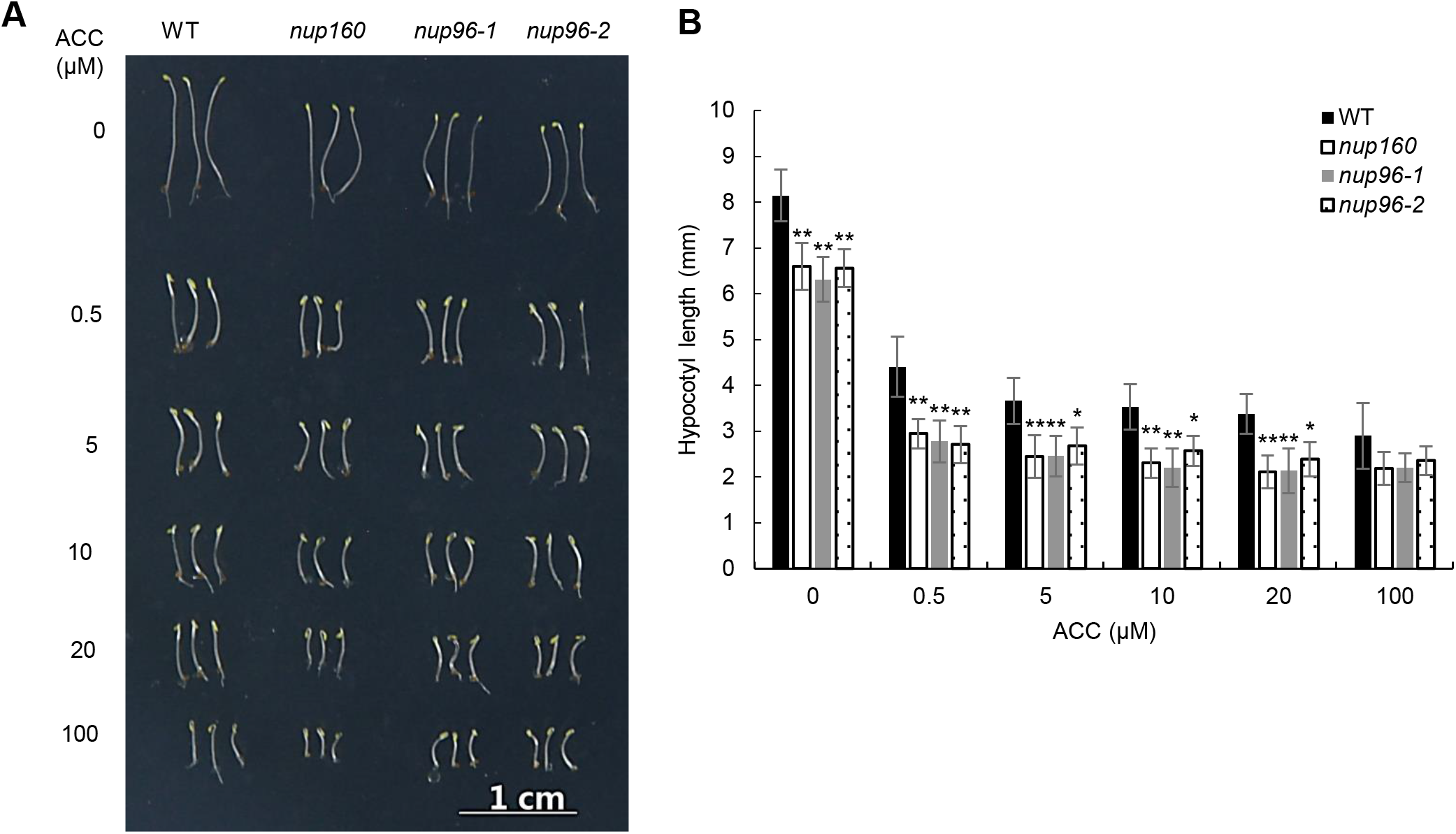
Ethylene “triple response” assays in nucleoporin mutants. (A) Ethylene sensitivity analysis of the wild type (WT) and nucleoporin mutants treated with or without ACC for 4 d. Bar= 1 cm. (B) Quantitative analysis of hypocotyl lengths for the WT and nucleoporin mutants grown on 1/2 MS medium containing different concentrations of ACC for 4 d under dark conditions. Values are mean ± SD; *p<0.05, **p<0.01.

As the expressions of the downstream ethylene-induced *ERFs* could be used as a reference for ethylene sensitivity as described previously (Wang *et al.*, 2017), we analyzed the relative expression levels of the ethylene-induced *ERFs* (*ERF1*, *ERF2*, *ERF4*, *ERF5*, *ERF11* and *ERF105*) in WT, *nup160*, *nup96-1* and *nup96-2*. Without ACC treatment, the expression levels of all the *ERFs* in WT and the nuleoporin mutants were low. Whereas, after ACC treatment, the expression levels of all the examined *ERFs* were promoted more in the mutants than in WT (Fig. 9), being consistent with the enhanced ethylene sensitivity of the nucleoporin mutants.

**Fig. 9.**
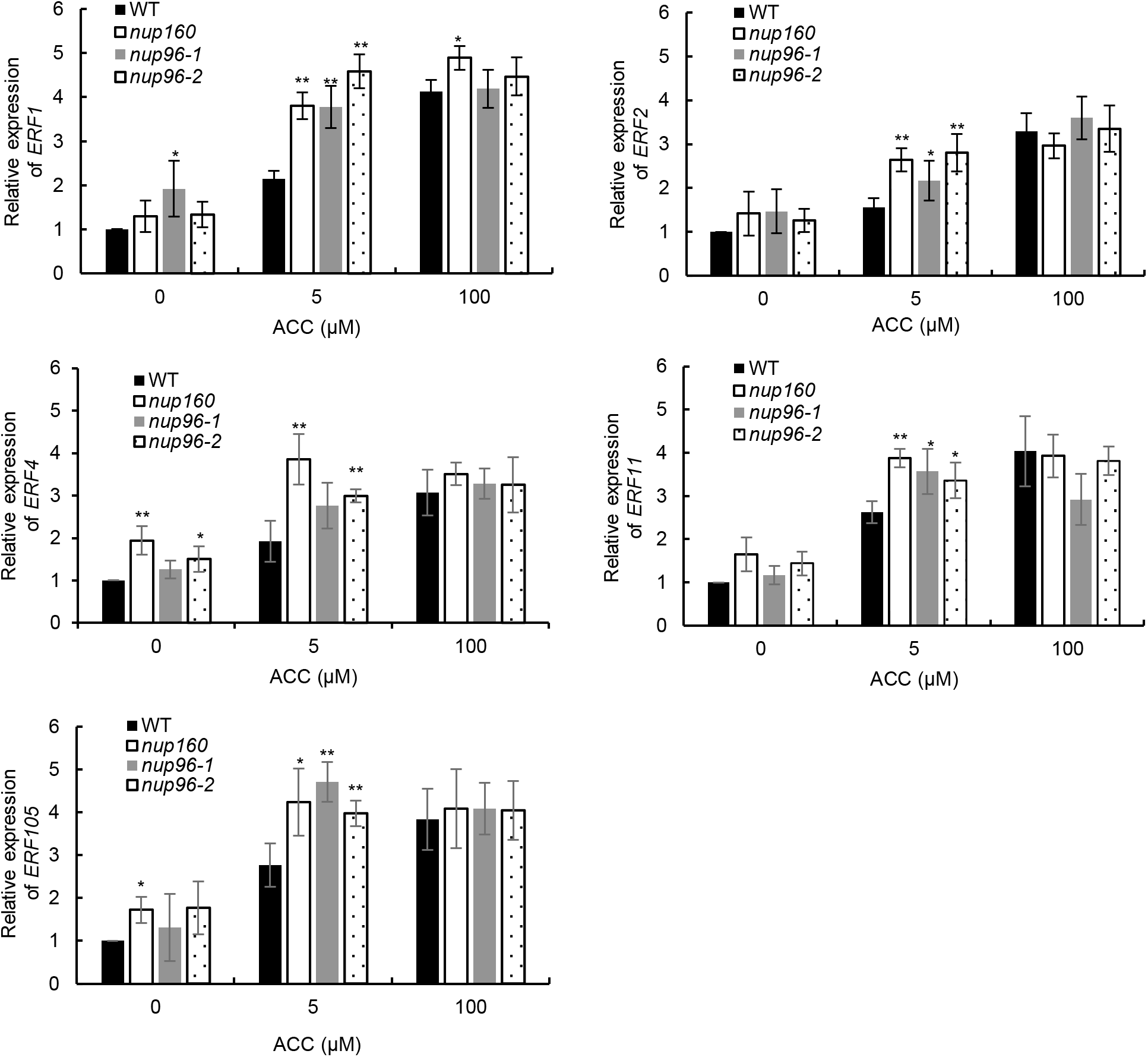
Relative expression of Arabidopsis *ERFs* in WT, *nup160, nup96-1* and *nup96-2*. The 10-day-old seedlings were grown on the medium with different concentrations of ACC for 6h. Values are mean ± SD; *p<0.05, **p<0.01.

## Discussion

The Arabidopsis *etr1-2* encodes an Ala^102^-to-Thr substitution in the ethylene binding domain, while the *etr1-1* contains a Cys^65^-to-Tyr substitution which is believed to prevent the copper binding and fully blocks ethylene binding (Schaller and Bleecker, 1995; Hall *et al.*, 1999; Rodriguez *et al.*, 1999). Previous study showed that the *rte1* mutations could restore the ethylene sensitivity of *etr1-2* but not *etr1-1* (Resnick *et al*., 2006), whereas the *cpr5* mutations could restore the ethylene sensitivity of both *etr1-2* and *etr1-1* (Wang *et al*., 2017), indicating that CPR5 functions differently from RTE1 in regulating the ETR1 receptor signaling. In this study, several appproaches were applied to investigate how CPR5 differs from RTE1 in the regulation, and at least three observations were obtained: (1) CPR5 binds to a larger area of the ETR1 N-terminus and needs at least three transmembrane domains for the binding, while RTE1 binds to a smaller region and needs only two transmembrane domains for the binding (Fig. 1); (2) Knockout of *CPR5* lead to a more enhanced ethylene sensitivity than that of *RTE1* (Fig. 3); (3) CPR5 acts as a nucleoporin in controlling the nucleocytoplasmic transport of mRNAs in ethyline signaling pathway, whereas RTE1 does not (Fig. 4; Fig. 5). These observations significantly advanced our understanding of the regulatory mechanism of CPR5 in ethylene signaling pathway.

RTE1 is mainly localized in ER and Golgi and required for the function of ethylene receptor ETR1 (Resnick *et al.*, 2006; Zhou *et al.*, 2007; Dong *et al.*, 2008). Genetic analysis revealed that Arabidopsis *RTE1* is required for the *ETR1* receptor signaling, but is not required for function of the other ethylene receptors, suggesting that RTE1 plays a specific role in regulating the ETR1 signaling (Resnick *et al.*, 2006). Similar genetic analysis using the double mutants of *cpr5-T3* with each of the ethylene insensitive mutants (*etr1-2, etr1-1, ers1-10, etr2-1, ein2-1*) showed that the ethylene insensitivity of the *ETR1* mutants (*etr1-2, etr1-1*) was suppressed by *cpr5-T3*, but not the others (*ers1-10, etr2-1, ein2-1*) (Wang *et al.*, 2017), suggesting that alteration of the ethylene sensivity of the mutants could not be totaly attributed to the defect of mRNAs nucleocytoplasmic transport as detected in fig. 5. The ETR1 ethylene receptor is the main target of CPR5 in ethylene signaling pathway. As an ETR1 associated protein, CPR5 could regulate the ETR1 receptor signaling through protein-protein interaction with the receptor. As supported, both CPR5 and RTE1 proteins are localized in ER and Golgi (Gu *et al.*, 2016; Wang *et al.*, 2017), and they could interact directly (Wang *et al.*, 2017). By yeast split-ubiquitin assay and BiFC experiment, it was showed that both CPR5 and RTE1 could bind to the N-terminus of ETR1 (1-368 aa) (Fig. 1). Further precise analysis indicated that RTE1 could interact with ETR1 (1-78 aa), while CPR5 did not (Fig. 1). CPR5 needs at least three transmembrane domains of ETR1 for the binding, while RTE1 needs only two transmembrane domains for the binding. The different requirement for the ETR1 domain binding between CPR5 and RTE1 might be one of the reasons why RTE1 differs from CPR5 in regulating the ETR1 receptor signaling. As the binding sites were generally the same, it was speculated that there might exist a competitive binding between CPR5 and RTE1 when they interacted with the ETR1 receptor.

It is worth to note that the single mutant *cpr5-T3* showed more enhanced ethylene response than *rte1-3* (Fig. 3). In addition, it was observed that the hypocotyl length of *cpr5-T3 rte1-3* was closer to that of *cpr5-T3* but not *rte1-3* when treated with different concentrations of ACC, suggesting that CPR5 might act downstream of RTE1 and play a more general regulatory function in ethylene signaling. As supported, this study provided evidence demonstrating that the nucleocytoplasmic transport of the *ETR1* mRNA and the other ethylene signaling related mRNAs (*ETR2*, *ERS1*, *ERS2*, *EIN4*, *CTR1*) was restricted in *cpr5-T3* but not in *rte1-3* or *rth-1* (Figs. 4, 5).

CPR5 is originally isolated from the research on plant pathogenesis (Bowling *et al.*, 1997; Boch *et al.*, 1998). Arabidopsis CPR5 has five transmembrane domains at its C-terminus and nuclear localization signal at the N-terminus. Subcellular localization analysis indicated that CPR5 was localized in the ER and Golgi membrane, and it was also localized in the nuclear membrane (Gu *et al.*, 2016). CPR5 was reported to be able to act as a nucleoporin in controlling of triggering immunity and programmed cell death (Gu *et al.*, 2016). In this study, we used poly(A)-mRNA *in situ* hybridization to examine whether CPR5 was invovled in the nucleocytoplasmic transport of mRNAs, the results showed that there was obvious mRNAs accumulation in the nucleus of *cpr5-T3* (Fig. 4). Further quantification analysis revealed the accumulation of *ETR1* mRNA in the nucleus of *cpr5-T3* cells accounted for more than 60% of its total level, while that in the nucleus of WT, *rte1-3* and *rth-1* was only about 30% (Fig. 5). We also analyzed the mRNAs accumulation of the other ethylene receptor encoding genes *ETR2*, *ERS1*, *ERS2* and *EIN4*, the downstream regulator encoding genes *CTR1*, *EIN2*, *EIN3*, and some ethylene-induced *ERFs* in the nucleus of *cpr5-T3*. The results showed that mRNAs of the *ETR2*, *ERS1*, *ERS2*, *EIN4* and *CTR1* were remarkably accmulated in the nucleus of *cpr5-T3*. In contrast, the mRNAs of the other detected genes were not dramatically accumulated in the nucleus of *cpr5-T3*. These results suggested that CPR5 could selectively control the nucleocytoplasmic transport of mRNAs in ethylene signaling pathway. Unfortunately, how these mRNAs are identified and controlled for the nucleocytoplasmic transport has not been understood yet.

In order to know whether CPR5 acts the same as the other nucleoporins in controlling the nucleocytoplasmic transport of mRNAs, we investigated mRNAs accumulation in the nucleoporin mutants *nup160*, *nup96-1* and *nup96-2* (Figs. 6, 7). The results showed that all the examined ethylene signaling pathway related mRNAs, including *ERT1, ETR2, ERS1, ERS2, EIN4*, the downstream *CTR1, EIN2, EIN3*, and the ethylene-induced *ERFs*, were accumulated in the nucleus of *nup160, nup96-1* and *nup96-2.* These observations indicate that CPR5 functions differently from the nucleoporin NUP160 or NUP96 in controlling the nucleocytoplasmic transport of mRNAs in ethylene signaling pathway.

Nucleoporin is a basic unit of the nuclear pore complex (NPC) which controls the two-way transport of RNA and protein between nucleus and cytoplasm. Yeast NPC contains 35 to 50 unique proteins (Yang *et al.*, 1998; Rout *et al.*, 2000), while mammalian NPC is a larger complex consisting of 80 to 100 unique proteins (Görlich and Kutay, 1999). In plants, at least 30 nucleoporins were identified, involved in different events including flowering, pathogen interaction, nodulation, cold response, and hormone signaling (Zhang and Li, 2005; Dong *et al.*, 2006; Parry *et al.*, 2006; Meier and Brkljacic, 2009; Tamura *et al.*, 2010). The NUP160 and NUP96 are the plant homologs of the vertebrate nucleoporins (Zhang and Li, 2005; Dong *et al.*, 2006), while CPR5 is a plant-specific transmembrane nucleoporin that may contribute to the stability of the NPC core scaffold (Gu *et al.*, 2016). Nuceoporins play multiple roles in various biological processes. However, the regulatory function of nucleoporin in plant response to ethylene has not been understood yet. In this study, we showed that the mRNAs of the ethylene related genes were accumulated in the nucleus of *nup160, nup96-1* and *nup96-2*, and finally led to enhanced ethylene sensitivity and elevated *ERFs* expression levels in the mutant plants (Figs. 8, 9).

In conclusion, this study investigated the binding sites of CPR5 and RTE1 to the ETR1 receptor and the results suggested that both of them could bind to the N-terminal transmembrane domains of ETR1. The regulation mode of CPR5 is different from that of RTE1 in ethylene signaling. CPR5 could regulate ETR1 by binding the ETR1 receptor and controlling its mRNA nucleocytoplasmic transport, while RTE1 regulates the ETR1 receptor signaling by a specific association with the receptor. In addition, it was found that the disruption of CPR5, NUP160 or NUP96 lead to the defective nucleocytoplasmic transport of ethylene related mRNAs. By comparing the effects of the nucleoporin proteins on the nucleocytoplasmic transport of mRNAs, we conclude that CPR5 might selectively control the nucleocytoplasmic transport of mRNAs, while NUP160 and NUP96 control bulk mRNA cytoplasmic export in ethylene signaling pathway. Although there are still some questions unanswered, the study provides a strong base for revealing the regulatory mechanism of ethylene receptor ETR1 and its regulatory factors.

## Supplementary data

Fig. S1. Relative hypocotyl growth of the WT and nucleoporin mutants.

Table S1. The primers used to construct yeast split-ubiquitin assay vectors

Table S2. Primers used for the constructs of the BiFC assays

Table 3. Primers used for qPCR

## Acknowledgements

We thank the Experimental Center of Qingdao Agricultural University for confocal microscopy support (Leica TCS SP5). This work was supported by the National Natural Science Foundation of China (#31870255; #31370322), the Shandong Natural Science Foundation (ZR2019MC061) and the Shandong Agricultural Variety Project (2019LZGC015) to CHD; and National Nature Science Foundation of China (#31600224) to Y.Y.

## Author contributions

J.C. performed most of the research and wrote the first draft of the manuscript. Y.Y., X.S. and L.Q. performed some yeast experiment and mutant screening. C.H.D. designed the experiments and revised the manuscript.

